# SERODIAGNOSIS OF SALMONELLA INFECTION: USING A LOGISTIC REGRESSION MODEL

**DOI:** 10.1101/2023.03.26.534120

**Authors:** James A Ndako, Iyanuoluwani J. Owolabi, Surajudeen A. Junaid, Victor T. Dojumo, Christy J. Ndako, Oluwakemi E. Abiodun, Remilekun M. Omoniyi, Sharon O. Bello

## Abstract

Salmonella infection remains a major global health problem and worsened by lack of appropriate diagnostic tools to aid early detection and teatment, particularly in low-income nations. *Salmonella typhi* is the most common causative agent of typhoid fever and the prevalence of this illness has been on the increase specifically in areas of poor personal hygiene and sanitation. This study was carried out to further improve the diagnosis of salmonella infection, through a mathematical regression model. An analysis was performed using the logistic regression approach and the predictability of the model was done by extracting fifteen (15) typhoid observations from the obtained samples; for the model to predict their status. The model was able to accurately predict 66.7% of the observations. This study showed an increased prevalence in typhoid fever including a significant correlation between typhoid fever and other parameters. The global burden of this illness can be minimized by proper vaccination, and prompt but appropriate diagnosis and treatment.Further studies and test-meaasures also needs to be carried out to improve diagnosis and treatment regimen.

## Introduction

Enteric fever is caused by *Salmonella enterica* serovar *Typhi* and also *Salmonella paratyphi*, which is a gram-negative rod-like shaped bacterium. This disease is prevalent in areas of low environmental sanitation and personal hygiene (1). Recently, the prevalence rate was recorded as 13 million cases affected annually and this is most common in both developing countries and undeveloped countries. The causative organism is a genus and consists of gram-negative, rod- shaped bacteria (2). They belong to the family Enterobacteriaceae and are intracellular pathogens, with over 2,300 serotypes. The as the typhoid-causing serotypes are only transferable from humans to humans (3). One of the main main sources of spreadof this agent is through the constant excretion of the bacterium in faeces of infected persons and recently-recovered individuals (4). These Samonellaserotypes are responsible for typhoid fever, paratyphoid fever and food borne infections (5). Invasion of the bloodstream by *Salmonella* causes typhoid fever and this can lead to the invasion of other organs in the body,(6). Symptoms of this illness inckudes headache, fever, diarrhea, abdominal pains among others after an incubation period of 1-2weeks (7). Proper and accurate diagnosis coupled with prompt treatment are effective means of avoiding further complications.

### Methodology

The data for this analysis was obtained was from the Landmark University Medical laboratory section. After proper ethical protocols and consent from the subjects;Two hundred (200) samples were obtained from the volunteer in and out patients of the Medical Facility. Widal test assay was carried out on the samples obtained, using appropriate test kit for widal analysis.Test were carried out based on the manufacturer′s instruction/maual.From the results of sera samples analysed and data obtained; A logistic linear regression was applied to the test data according to the model adopted by (8). This defines if or not a patient is typhoid fever positive towards the binary response variable, the model is designed to predict. Mathematically, the response variable which is the typhoid status of each patient was denoted as:

Y and represented as:

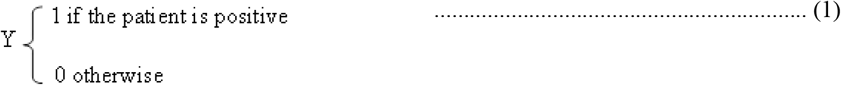

The logistic regression in terms of positive

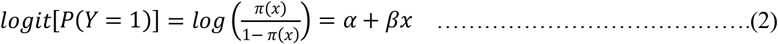

Where π(*x*) is equivalent to the probability of being positive, P(Y=1).

*x* contains the matrix of the explanatory variables.

The expression, (2) have a linear right-hand side and a logarithmic scale of the odds of being positive. With the exponentiation and algebraic manipulation, π(*x*) is given as (3) to obtain a form that results into values between 0 and 1 which are interpreted as probabilities.

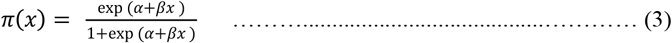

The typhoid model with respect to the variables included in this study is generally expressed as: Status ∼ f(Age-cat, PCV, WBC, NEUT, LYMP, MONO, PLT, HB, ESR) and mathematically as

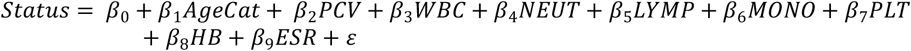

Where *β*_*i*_′*s* are the independent variables’ coefficients. The ages were categorized as 0 – 10, 11 – 20, 21 – 30, 31 – 40, 40 and above.

The Akaike Information Criterion (AIC); which is an estimator of prediction error and thereby relative quality of statistical models for a given set of data would be used to check the measure of fit. The AIC provides a measure of information that a model provides. It is used in measuring the tested model against the theoretically true model.

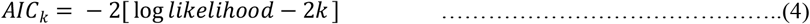

It is of interest to minimize (4) where k is the number of parameters in the model. Thus, for simplicity, independent variables are been reduced until the smallest AIC is obtained for a befitting model.

Other measures of fitness may also be considered in this process such as the Deviance, McFadden’s pseudo R^2^, concordance and the significance of model parameters. The deviance measures against the most complex model possible, a saturated model with an individual parameter for each observation. It follows an approximately chi-squared distribution and the deviance likelihood ratio statistic tests the hypothesis that all parameters not used in the tested model are zero. McFadden’s pseudo R^2^ is different from the common R^2^ because it does not compare variances but it also ranges from 0 to 1. The general impression is that a higher value is more desirable with ‘good’ fitting models in the 0.2 – 0.4 range(9). Concordance measures predictive power which is estimated from a receiver operating characteristic (ROC) curve. The curve is a plot of model sensitivity versus (1 – specificity). The sensitivity of a model can be defined as the probability that the model predicts positive given that it is negative (10). The total area under the curve (AUC) is equivalent to concordance. (8) value of AUC = 0.5 is equivalent to guessing. Thus, a value closer to one is the better. The independent variables were also described through the use of statistics such as the bi-variate correlation coefficients. The accuracy of the model is also tested by using extracted typhoid patients’ data

## Results and Discussion

The combined paired histograms, scatter plots and correlation coefficients is presented in a square matrix (Figure 1). The display helps to visualize how the continuous independent variables are related. The correlation coefficients are reported in the upper triangular portion of the matrix in Figure 1. The highest correlation was observed between PCV and HB with 0.98 which is strong positive and, NEUT and LYMP with −0.98 which is strong negative. WBC and NEUT, WBC and LYMP, NEUT and MONO also have a moderate correlation, 0.44, −0.43 and −0.42 respectively. The linearity of the relationship can also be observed from the scatter plot.

**Figure 1.**
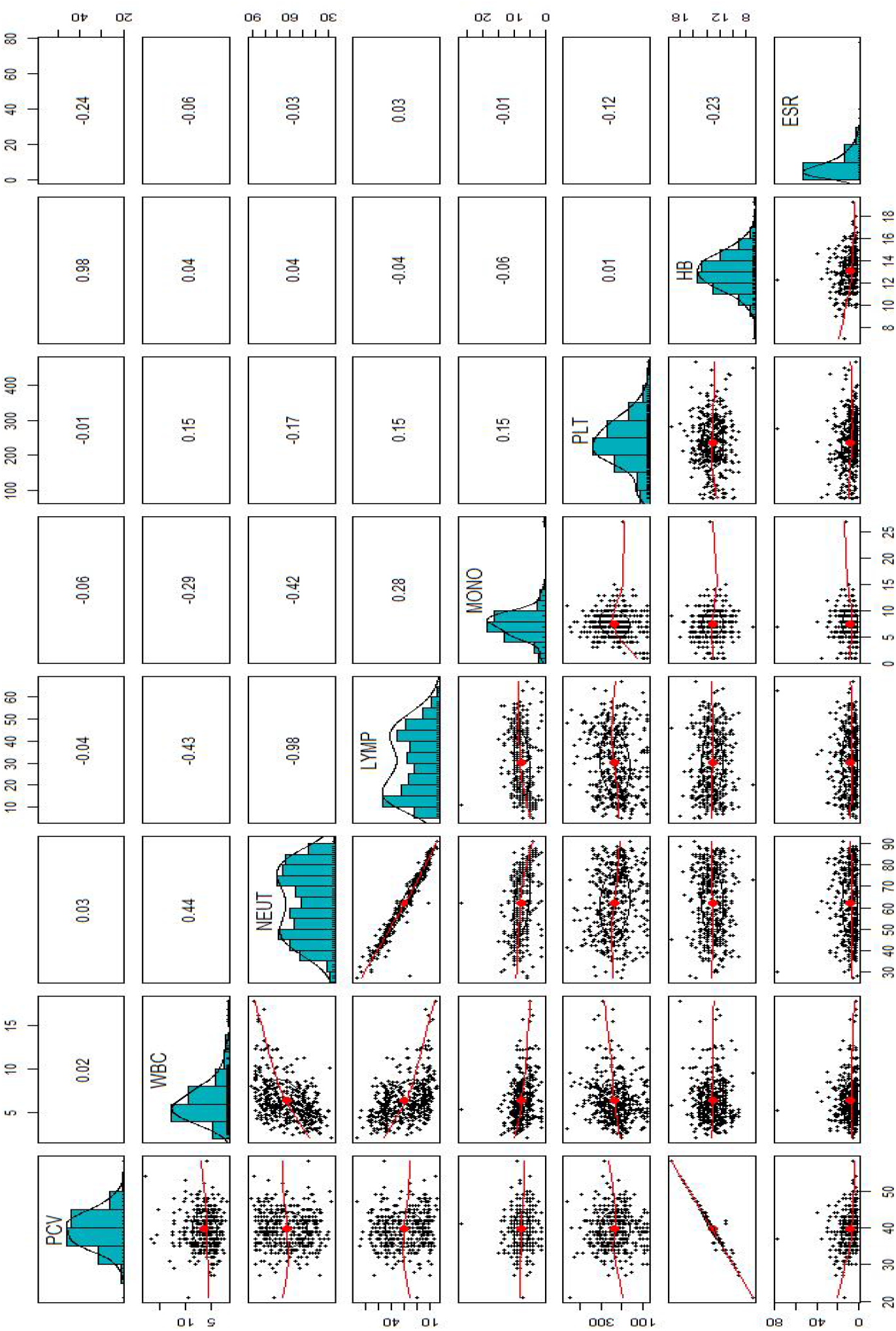
: Descriptive and Correlations of the Independent Variables.

Table 1 presents the possible model through removal of some independent variable and their respective AIC. Model 1 is the originally fitted model which has an AIC of 422.95 and while Model 6 which has WBC, PLT, HB and ESR as its independent variable has the least AIC value. Thus, the simplest form of Model 1 with the most relevant variables as Model 6. The logistic regression handles a categorical independent variable such as age category in a different manner; it takes the first category as a base and assigns binary figures to the rest of the category just like the case of a dummy regression. The age categories, PCV, NEUT, LYMP, MONO and HB were not statistically significant in model 1 at p-value > 0.05. A unit increase in WBC and PLT will significantly reduce the log odds by 0.1372 and 0.0112 respectively while a unit increase in ESR will increase the log odds by 0.1430 (Table 2).

**Table 1:** Most fitted model through the AIC Criterion.

| MODEL | Included Variables | AIC |
| --- | --- | --- |
| 1 | Status ~ Age-cat + PCV + WBC + NEUT + LYMP + MONO + PLT + HB + ESR | 422.95 |
| 2 | Status ~ PCV + WBC + NEUT + LYMP + MONO + PLT + HB + ESR | 419.93 |
| 3 | Status ~ PCV + WBC + LYMP + MONO + PLT + HB + ESR | 418.04 |
| 4 | Status ~ PCV + WBC + LYMP + PLT + HB + ESR | 416.11 |
| 5 | Status ~ WBC + LYMP + PLT + HB + ESR | 414.24 |
| 6 | Status ~ WBC + PLT + HB + ESR | 413.06 |

**Table 2:** Logistic Regression Coefficient Result for Model 1.

| **Coefficients** | **Estimate** | **Std. Error** | **Z value** | **P-value** |
| --- | --- | --- | --- | --- |
| Intercept | 13.0696 | 8.0051 | 1.633 | 0.1025 |
| Age Category 2 | -0.9087 | 0.8595 | -1.057 | 0.2904 |
| Age Category 3 | -0.6232 | 0.8675 | -0.718 | 0.4725 |
| Age Category 4 | -0.4527 | 1.0050 | -0.450 | 0.6524 |
| Age Category 5 | 0.8829 | 1.4255 | 0.619 | 0.5357 |
| PCV | -0.0344 | 0.1213 | -0.284 | 0.7765 |
| WBC | -0.1372 | 0.0649 | -2.113 | 0.0346 |
| NEUT | -0.0352 | 0.0774 | -0.455 | 0.6489 |
| LYMP | -0.0456 | 0.0783 | -0.582 | 0.5603 |
| MONO | -0.0490 | 0.0917 | -0.534 | 0.5935 |
| PLT | -0.0112 | 0.0021 | -5.391 | $7.01 \times 10^{-8}$ |
| HB | -0.3547 | 0.3553 | -0.998 | 0.3181 |
| ESR | 0.1430 | 0.0250 | 5.723 | $1.05 \times 10^{-8}$ |

The simplified model presents the model intercept, PLT, HB and ESR to be statistically significant at p-value < 0.001 while WBC can only be significant at p-value < 0.1. A unit increase in the WBC, PLT and HB reduces the log odds by 0.0955, 0.0118 and 0.4512 respectively while a unit increase in ESR will increase the log odds by 0.1449. Also, a very small p-value suggests a strong relationship between the typhoid status of patients and PLT, HB and ESR each (Table 3). The respective description of the deviance residuals for the two models (Model 1 and 6) are observed to be close as presented in Table 4. In model 1, adding PCV, WBC, NEUT, PLT and ESR significantly reduces the residual deviance at p-value < 0.05 while the other variables seem to improve the model less. A large p-value indicates that the model without the variable explains more or less the same amount of variation (Table 5). Adding WBC, PLT, HB and ESR in model 6 significantly reduces the residual deviance at p-value < 0.01 (Table 6). A good fit can be inferred from the McFadden values for both models (Table 7), since the value lies within the 0.2 – 0.4 range according to (McFadden, 1974). Figure 2 and 3 presents the receiver operating characteristic (ROC) curve for model 1 and 6 respectively. The area under the curve for model 1 and 6 was obtained to be 0.6428 and 0.6607 which is better because it is greater than 0.5, that is, more than guessing.

**Table 3:** Logistic Regression Coefficient Result for Model 6.

| **Coefficients** | **Estimate** | **Std. Error** | **Z value** | **P-value** |
| --- | --- | --- | --- | --- |
| Intercept | 8.1534 | 1.4094 | 5.785 | $7.25 \times 10^{-9}$ |
| WBC | -0.0955 | 0.0550 | -1.735 | 0.0827 |
| PLT | -0.0118 | 0.0020 | -5.918 | $3.25 \times 10^{-9}$ |
| HB | -0.4512 | 0.0872 | -5.172 | $2.32 \times 10^{-7}$ |
| ESR | 0.1449 | 0.0246 | 5.878 | $4.15 \times 10^{-9}$ |

**Table 4:** Descriptive of the Deviance Residuals.

| Model | Min | 1Q | Median | 3Q | Max |
| --- | --- | --- | --- | --- | --- |
| 1 | -4.5829 | --0.8514 | -0.0289 | 0.7900 | 2.7539 |
| 6 | -4.5062 | -0.8554 | -0.0352 | 0.7893 | 2.7323 |

**Table 5:** Analysis of Deviance for Model 1.

| Included Variables | df | Deviance | Residual df | Residual Deviance | P-value |
| --- | --- | --- | --- | --- | --- |
| NULL |  |  | 399 | 554.52 |  |
| Age Category | 1 | 8.946 | 395 | 545.57 | 0.0625 |
| PCV | 1 | 42.069 | 394 | 503.50 | $8.810 \times 10^{-11}$ |
| WBC | 1 | 8.931 | 393 | 494.57 | 0.0028 |
| NEUT | 1 | 9.026 | 392 | 485.55 | 0.0027 |
| LYMP | 1 | 0.024 | 391 | 485.52 | 0.8764 |
| MONO | 1 | 1.514 | 390 | 484.01 | 0.2185 |
| PLT | 1 | 42.417 | 389 | 441.59 | $7.375 \times 10^{-11}$ |
| HB | 1 | 1.060 | 388 | 440.53 | 0.3033 |
| ESR | 1 | 43.576 | 387 | 396.95 | $4.077 \times 10^{-8}$ |

**Table 6:** Analysis of Deviance for Model 6.

| Included Variables | df | Deviance | Residual df | Residual Deviance | P-value |
| --- | --- | --- | --- | --- | --- |
| NULL |  |  | 399 | 554.52 |  |
| WBC | 1 | 8.651 | 398 | 545.87 | 0.0033 |
| PLT | 1 | 43.834 | 397 | 502.03 | $3.575 \times 10^{-11}$ |
| HB | 1 | 52.121 | 396 | 449.91 | $5.217 \times 10^{-13}$ |
| ESR | 1 | 46.851 | 395 | 403.06 | $7.661 \times 10^{-12}$ |

**Table 7:** Measure of Fitness – McFadden’s pseudo R2.

| Model | llh | llhNull | G2 | McFadden | r2ML | r2CU |
| --- | --- | --- | --- | --- | --- | --- |
| 1 | -198.48 | -277.26 | 157.56 | 0.2841 | 0.3256 | 0.4341 |
| 6 | -201.53 | -277.26 | 151.46 | 0.2731 | 0.3152 | 0.4202 |

**Figure 2:**
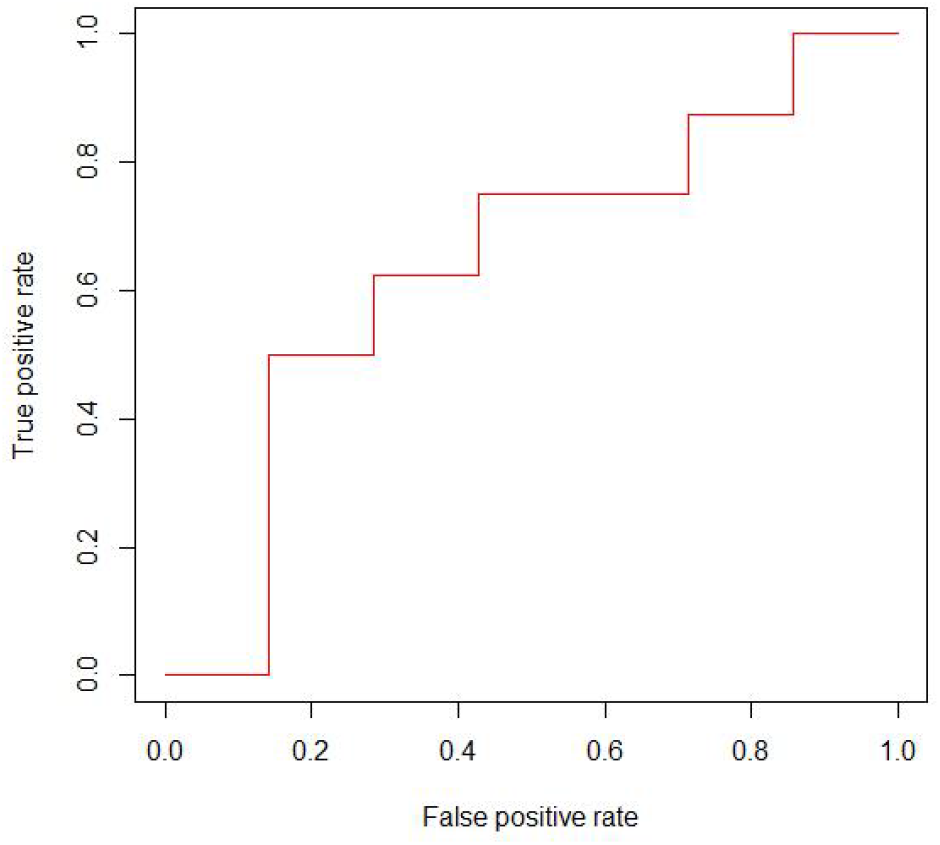
ROC Curve for Logistic Model 1.

**Figure 3:**
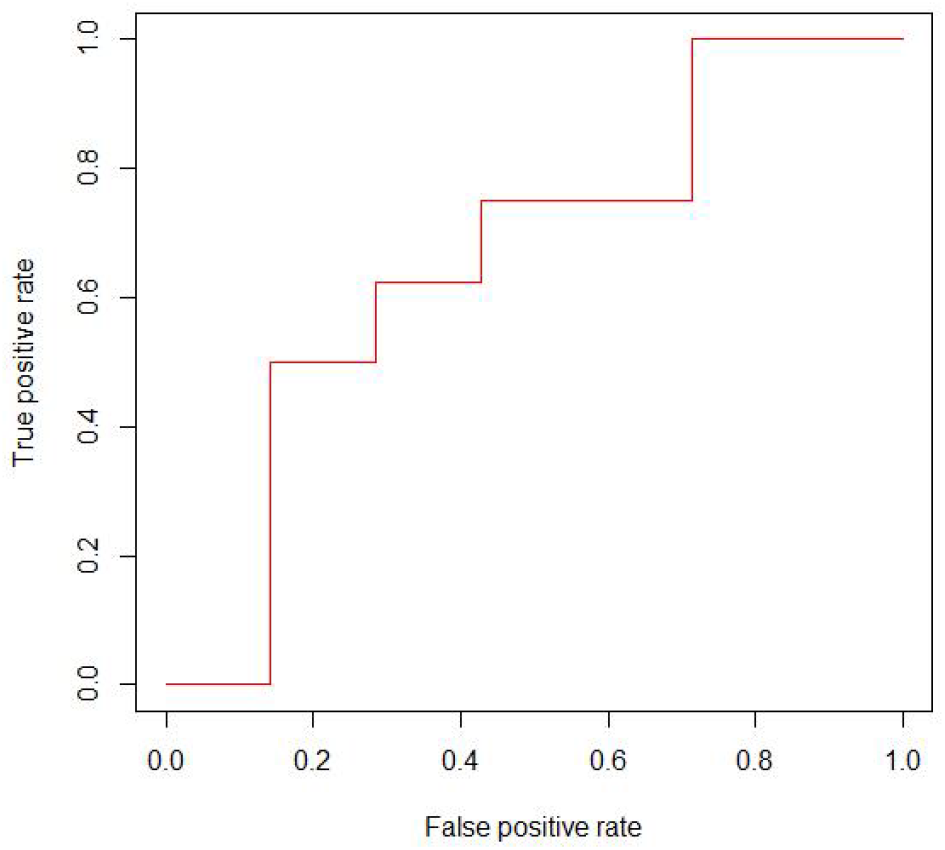
ROC Curve for Logistic Model 6.

### Assessing the predictive ability of the model

To further assess the predictability of the model, fifteen (15) typhoid observations from the positive sera samples were extracted for the model to predict their status. The model was able to accurately predict 66.7% of the observations,which could further validate the accuracy of diagnosis.This predictability status would undoubtedly aid prompt diagnosis and limit the transmission of the infectious agent.

## Conclusion

Studies carried out on typhoid and paratyphoid infection have emphasized the increasing prevalence among the population.This observation showed that a significant correlation exists between typhoid fever and hematological changes with the logistic regression approach employed.As a safeguard to further spread, better awareness, prompt diagnosis and vaccination are efficient measures that could be adopted to reduce the global burden of this infection.

## Notes

### Competing Interest Statement

The authors have declared no competing interest.

